# GDOP: A graph convolutional network-based drug “on-target” pathway prediction algorithm

**DOI:** 10.1101/2024.03.03.583216

**Authors:** Xiaolong Wu, Lehan Zhang, Mingyue Zheng

## Abstract

Since most compounds do not induce changes in the transcriptomic levels of their target proteins in vivo, traditional gene set enrichment analysis methods can only retrieve downstream differentially expressed genes, which offer little hints to their targets. To address this problem, we proposed a graph convolutional network-based drug “on-target” pathway prediction algorithm, GDOP, which can predict small pathways that contain target gene through the power of deep learning algorithms. Our model receives as input structural information and biological characteristics (gene expression profiles) of molecules. After being trained on the publicly available LINCS data set, GDOP showed better generalization ability, reaching an AUC-ROC of 0.89 and an averaged Top10 accuracy of 0.63 on the test set. Besides, demonstrated that GDOP was able to use RNA-Seq data as input and achieved accuracy prediction results.

## 1. Introduction

Understanding the mechanism of action (MoA) of biologically active compounds is a pivotal step in early drug discovery and development. Biologically active compounds exert their MoA phenotypes by interacting with their in vivo targets, causing in vivo omics (i.e., transcriptome) level changes. As a result, the development of omics technology and curated biological pathways provide a new perspective for presuming MoA of candidate compounds^1–5^.

Traditional methods typically use chemically inducible changes in transcriptional profiles and gene set enrichment analysis (GSEA) to evaluate MoA of chemical molecules^1, 6–8^. They seek for mathematically significant pathways by perform GSEA on compound-induced signatures. However, most of these mathematically significant pathways are “off-target” pathways, as the target genes of molecules are not settled in the pathways. This is due to the fact that the target genes of most molecules do not embed in the differentially expressed genes (DEG)^9^. At a result, the “off-target” pathways are only marginally helpful to identify the targets of the biologically active compounds.

The latest progress in the application of deep learning in the field of life sciences shows the unparalleled charm of artificial intelligence algorithms. Deep learning algorithms as enabling tools are used to help with molecular encoding, chemical synthesis route planning and inhibitor target prediction^10–13^. We envisioned that a model capable of predicting “on-target” pathways for new compounds with potential biological activity would make it much easier to find out targets of the compounds. The “on-target” pathways, as opposed to “off-target” pathways, refer to small pathways with at least one target gene as a member inside. It is not difficult to find that using pathways rather than target genes as training tasks can enhance the predictive power of those targets with few ligands. First, both simplified molecular-input line-entry system (SMILES) chemical encoding and compound-induced signatures that were measured in the L1000 project are used as inputs to fit “on-target” pathways in our developed neural network^14^. Our model was referred as graph convolutional network-based drug “on-target” pathway prediction algorithm (GDOP).

## 2. Results

### 2.1 The architecture and training of GDOP

The compound-induced signatures with PPI network were passed to a sGCN module to get latent vectors. The SMILES of chemical compounds were encoded into Morgan fingerprints. Then the latent vectors and the Morgan fingerprints were concatenated by the same compound and the united vectors were further fed to a deep dense network to predict the “on-target” pathways. (see Methods for details of all layers).

To take full advantage of the properties of molecules, both structural and biological features of molecules were used. Our previously published spectral-based graph convolutional network (sGCN) was applied to retrieve latent vectors from compound-induced signatures^13^ (Fig. 1, upper left). The sGCN is capable of unifying information on the topology of the PPI network and the gene expression profiles. The SMILES of chemical compounds were encoded into Morgan fingerprints using rdkit (version 2022.03.5)^15^ (Fig. 1, lower left). The latent vectors and the Morgan fingerprints were concatenated by the same compound and the united vectors were further fed to a deep dense network to predict the “on-target” pathways (Fig. 1, right).

**Fig. 1:**
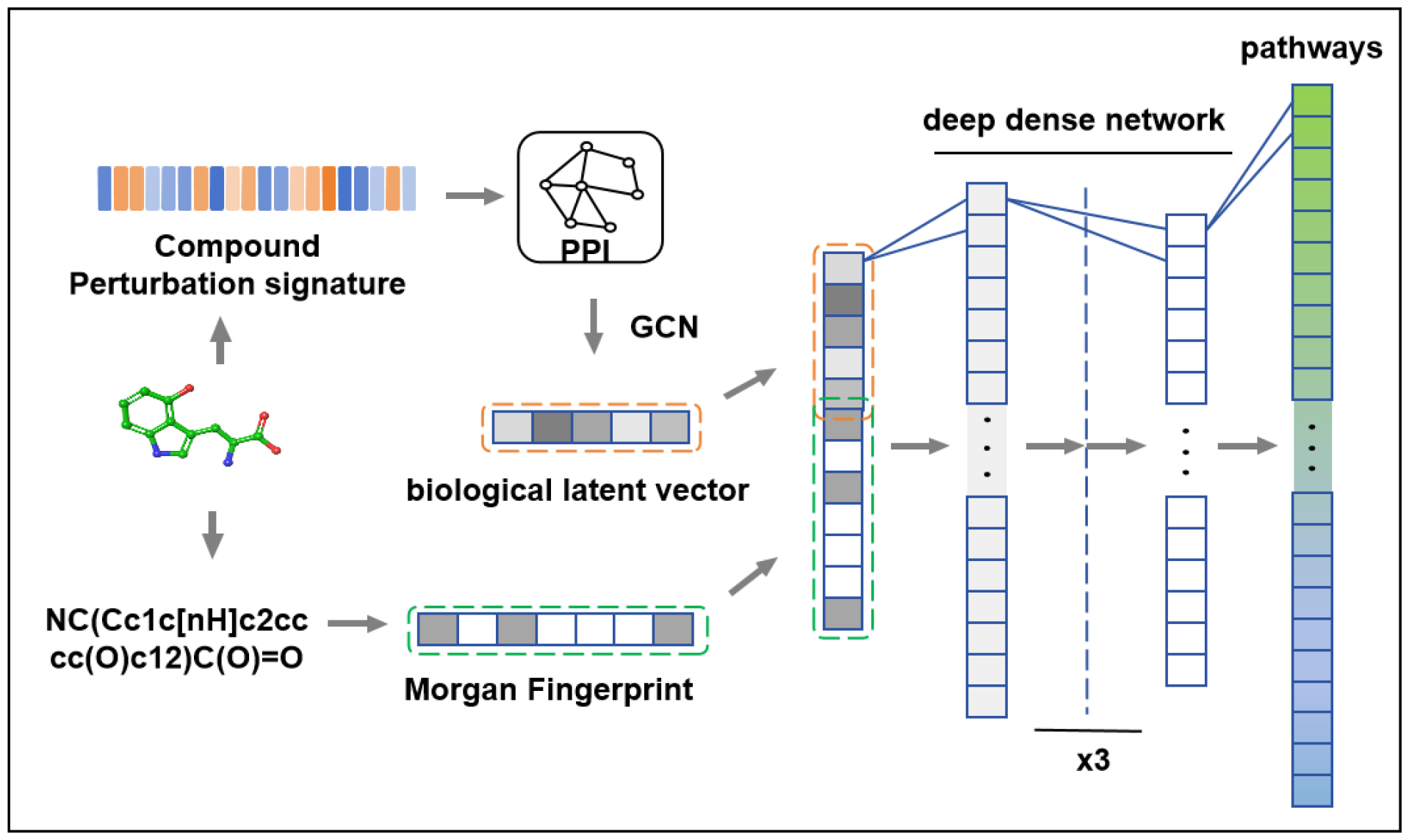
A schematic representation of the deep learning model underlying GDOP.

### 2.2 Model Training

The high-throughput gene expression profile data from the Library of Integrated Network-Based Cellular Signatures (LINCS) program was retrieved to fit our model. The LINCS dataset covers transcriptional changes induced by ∼2,400 small molecules with target information^14^. The dataset was updated by the latest LINCS Data Portal 2.0^16^. After cleaning, 2,348 molecules were randomly split to generate a training set (1,870), a validation set (234) and a test set (234). The candidate pathways were curated from the Human Molecular Signatures Database (MSigDB, version 2022.1)^17, 18^. We filter out relatively large pathways (pathways with more than 65 genes) and some pathways that are not related to the mechanism of action (for example, CYTOCHROME_P450_ARRANGED_BY_SUBSTRATE_TYPE). Finally, 1001 candidate target pathways were retained. The positive “on-target” pathways were labelled for each molecule according to its targets. Like in our previous research., for each compound three negative pathways were generated for each positive pathway through a random cross combination of compounds and pathways^13^.

By using the structural and biological features of the molecule and setting appropriate hyperparameters (see Methods), the model converges rapidly during training, as shown in Fig. 2A. The areas under the curve (AUC) of the receiver operating characteristic (ROC) for training, validation and testing are 0.99, 0.92 and 0.89 respectively, as shown in Fig. 2B.

**Fig. 2:**
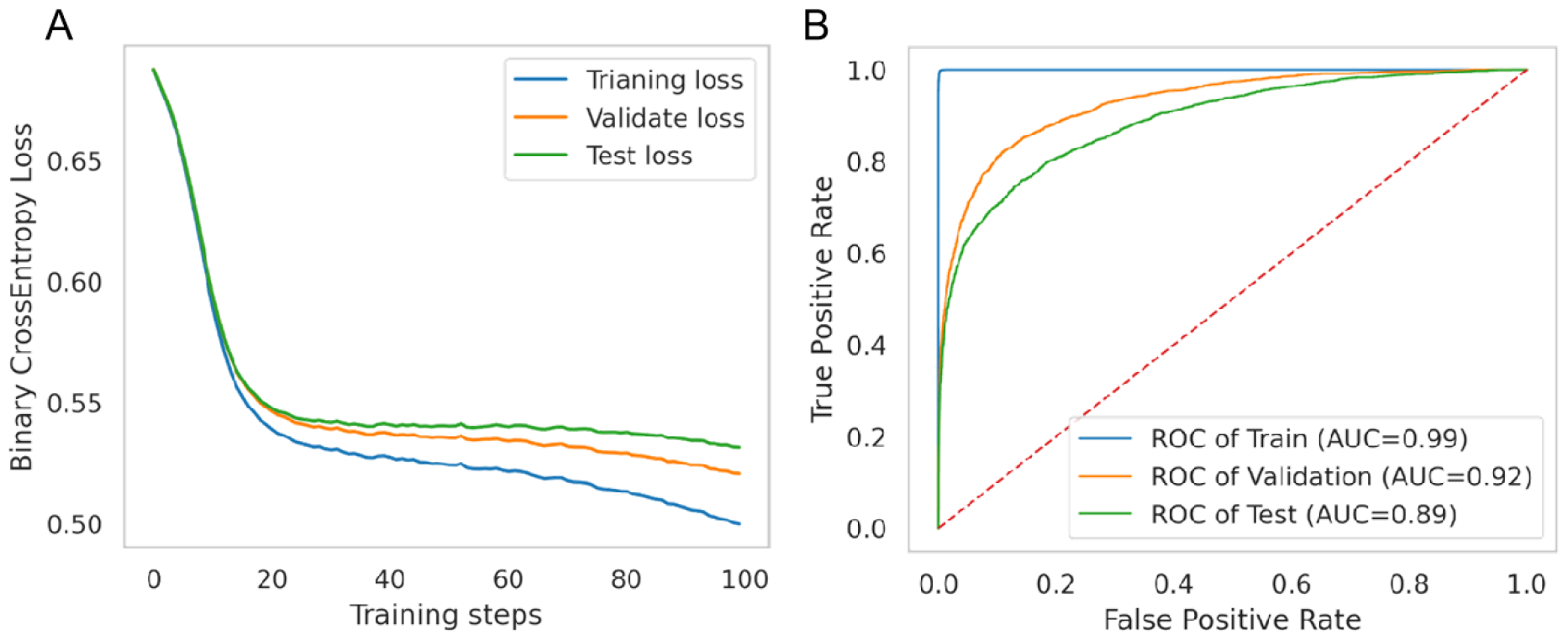
Training results of the GDOP.

A, The binary cross entropy loss of training, validation and test loss in the first 100 steps. B, The ROC curves of the training, validation and test dataset.

For ease of comparison, the same performance metric, Top N accuracy (N=10, 30 or 100), was used to evaluate the performance of our model (refer to Methods). As shown in Fig. 3, the top10 accuracy, the top30 accuracy and the top100 accuracy of the traditional method, single sample Gene Set Enrichment Analysis (ssGSEA), on the test set were on par with blind guessing (the random method)^6, 19^. As expected, our model GDOP achieves better performance on the corresponding test set. The average values of GDOP’s Top10 accuracy, Top30 accuracy, and Top100 accuracy are 0.63 (±0.02), 0.76 (±0.02), and 0.90 (±0.01) respectively.

**Fig. 3:**
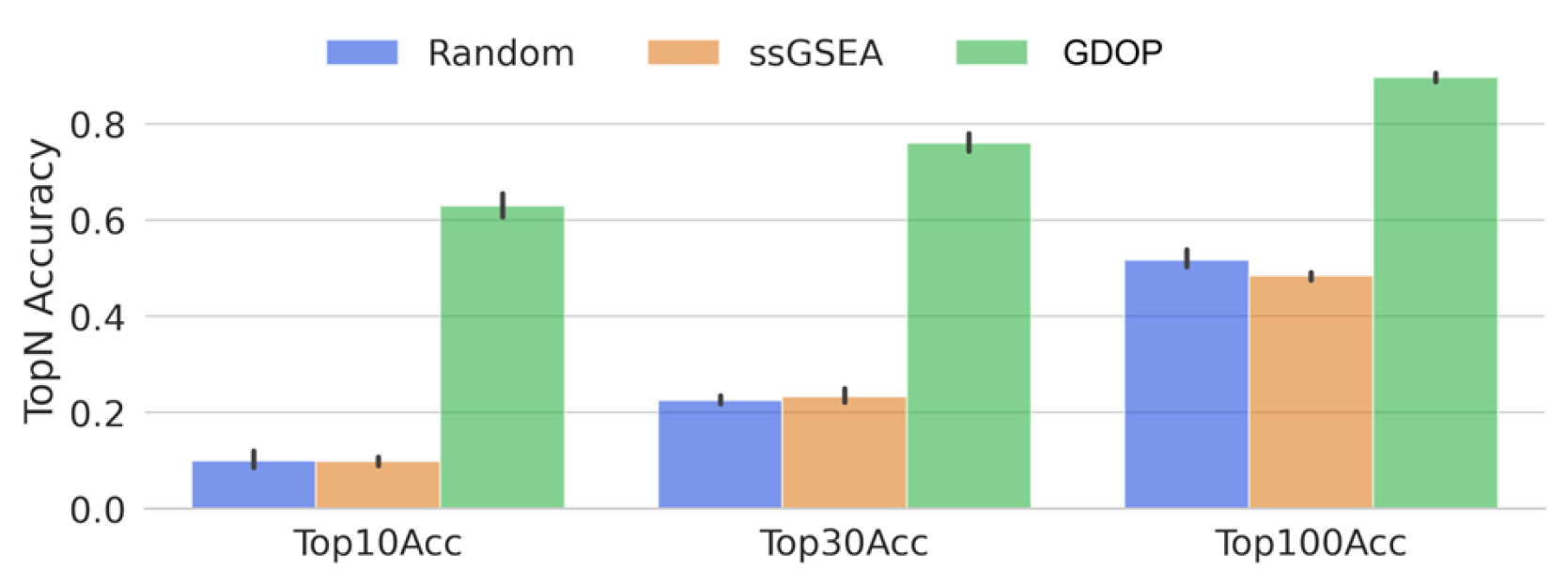
The top N (N=10, 30 or 100) accuracy on test set for different methods.

### 2.3 Ablation studies

Table 1 presents the results of the ablation studies to better understand GDOP prediction models. We varied the components of the base model and measured the classification performance on the validation set. In row (A) and (B), we can observe that disregarding the structure inputs or the biological inputs can lead to the degrade of the model performance. And the model performance relied more on structure information than biological information. In row (C), we can observe that disregarding the GCN module significantly degrades the model performance. This suggests that the GCN module can help to retrieve more useful latent vector from raw gene signature inputs.

**Table 1.** Ablation studies on the prediction model of GDOP.

|  | Input Features |  | GCN | Top10 Accuracy |
| --- | --- | --- | --- | --- |
|  | Perturbation signature | Compound structure |  |  |
| (A) | TRUE | FALSE | TRUE | 0.37 |
| (B) | FALSE | TRUE | FALSE | 0.63 |
| (C) | TRUE | TRUE | FALSE | 0.57 |
| BASE | TRUE | TRUE | TRUE | 0.70 |

### 2.4 Model application

To test whether the model can use data from RNA-Seq quantitative technology that is different from L1000 quantitative technology as input and make accurate prediction results, we designed a cell experiment as shown in Table 2. The processed biological samples were sent to Meji Biotechnology Company for on-machine sequencing using the Illumina NovaSeq Xplus platform. After sequencing, the data were quality controlled and filtered using fastp software, compared with HiSat2 software, and finally quantified using RSEM software ^20–22^. The version of the human reference genome used for alignment was GRCh38, and the raw read counts of each gene were obtained after quantification. Then the raw read counts were converted into model input using the method mentioned in section 4.5

**Table 2.** The cell experiment of s-ketamine.

| Compound | Cell line | Experiment condition | Sample number | Group |
| --- | --- | --- | --- | --- |
| s-ketamine | A549、MCF7 | 10um, 24h | 3+3 | trt |
| DMSO | A549、MCF7 | 10um, 24h | 3+3 | ctrl |

s-ketamine has rapid and long-lasting antidepressant effects. In 2019, the FDA approved it for the treatment of patients with treatment-resistant depression T (Treatment-Resistant Depression, referred to as TRD) and major depressive disorder (MDD) with suicidal ideation. Research shows that the main target of s-ketamine is the glutamatergic NMDAR (N-methyl-D-aspartate receptor)^23^. The prediction results of our proposed GDOP model for s-ketamine are shown in Table 3. Among them, the top three predicted pathways all contain NMDAR, the reported target of s-ketamine. The results demonstrated that GDOP was able to use RNA-Seq data as input and achieved accuracy prediction results.

**Table 3.** Top 10 candidate “on-target” pathways of s-ketamine predicted by GDOP.

| Ranking | Biological pathways in Reactome |
| --- | --- |
| 1 | Ras activation upon Ca<sup>2+</sup> influx through **NMDA** receptor |
| 2 | Unblocking of **NMDA** receptors, glutamate binding and activation |
| 3 | Assembly and cell surface presentation of **NMDA** receptors |
| 4 | GABA receptor activation |
| 5 | Negative regulation of **NMDA** receptor-mediated neuronal transmission |
| 6 | EPHB-mediated forward signaling |
| 7 | Synaptic adhesion-like molecules |
| 8 | Synthesis, secretion, and deacylation of Ghrelin |
| 9 | Acetylcholine binding and downstream events |
| 10 | Long-term potentiation |

## 3. Discussion

Omics quantitative technology, especially the cheap transcriptomics, can provide a comprehensive picture of certain microscopic molecules in cells, and has increasingly become a means to quickly evaluate the intracellular action mechanism and metabolism of new compounds. Since most compounds do not induce changes in the transcriptomic levels of their target proteins in vivo, traditional gene set enrichment analysis methods can only retrieve downstream differentially expressed genes, which offer little hints to their targets. To address this problem, we built a new model, GDOP, which can predict “on-target” pathways through the power of deep learning algorithms. Our model receives as input structural information and biological characteristics (gene expression profiles) of molecules. After being trained on the publicly available LINCS data set, GDOP showed better generalization ability, reaching an AUC-ROC of 0.89 and an averaged Top10 accuracy of 0.63 on the test set.

In addition, through practice, we found that GDOP can not be limited to the quantitative data of the L1000 platform. Illumina high-throughput transcriptome data can also be used as input to the GDOP model after undergoing the conversion process shown in section 4.5. As shown in section 2.4, by designing cell experiments for s-ketamine, we have completely verified the operability and practicality of the model to predict the target of the compound.

## 4. Method

### 4.1 Data collection

The LINCS phase I L1000 dataset (GSE92742) and updated version were downloaded from the Gene Expression Omnibus (GEO) and LINCS Data Portal 2.0, respectively^16, 24^. Apart from the signatures, the meta information including targets was curated from the source. The reactome pathways were obtained from the Human Molecular Signatures Database (MSigDB, version 2022.1)^18^. The human PPI network from the STRING (version 11.5) database was downloaded^25^.

### 4.2 Data processing

#### Reactome

The large pathways (i.e., over 65 genes) and certain MoA-unrelated pathways (i.e., CYTOCHROME_P450_ARRANGED_BY_SUBSTRATE_TYPE) are excluded, since large pathways are related to general biological pathways, rather than specific to a certain function^26^. Finally, 1001 curated reactome pathways were obtained.

#### LINCS

We filtered out the molecules: lack of target information, less five signatures or SMILES couldn’t be successfully parsed using rdkit (version 2022.03.5). The profiles for each molecule were averaged by ignoring plate, dose, treatment time and cell line details. We used only the landmark genes. Finally, 2,348 valid molecules were kept and randomly split into training (1,870), validation (234) and test (234) sets. The positive pathways for each molecule were labelled according to its target information using the curated reactome pathways mentioned above. Like in Zhong et al., for each compound three negative pathways were generated for each positive pathway through a random cross combination of compounds and pathways^13^.

#### STRING

We only kept the nodes present in the “landmark” gene set and the PPI edges with a “combined score” greater than or equal to 800. Accordingly, the curated PPI network consists of 978 nodes and 8,320 edges (including self-connection).

### 4.3 The training procedural

We applied both gene profile and structure features of molecules to train our GDOP model. The profile features were transformed into latent vectors by our early developed spectral-based GCN (sGCN) module in Zhong et al.^13^. The sGCN module helps to unify information on the topology of the PPI network and the differential gene expression profiles. The SMILES of chemical compounds were encoded into Morgan fingerprints using rdkit. Then we concatenated the latent vectors and the Morgan fingerprints and fed them into a feedforward neural network with four layers of 4096, 3016, 2048, 1024. The dropout rates for each layer, including sGCN layer, were 0.2, 0.5, 0.5, 0.5, 0.2. The rectified linear unit (ReLU) activation function was used for the output of each layer except for the last one. We used cross entropy as our cost function, and Adam as our optimizer algorithm. Early stopping was used to terminate the training process if the performance of the model on the validation dataset shows no further improvement in specified successive steps.

### 4.4 Model evaluation metric

We mainly used the same performance metric, top N accuracy, to evaluate the performance of our model and other methods. This metric reflects the proportion of tested compounds whose any true target can be correctly predicted among the top ranked N targets, and in this study, N values of 100, 30, 10 were evaluated.

### 4.5 Process of transformation from RNA-Seq raw read counts into model input

(1) Convert the count values 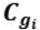 of each gene ***g**_i_* into log values 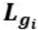:

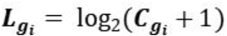

(2) Calculate the means 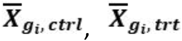 and variances 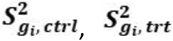 of each gene log values 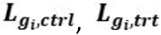 in control and compound treated groups respectively:

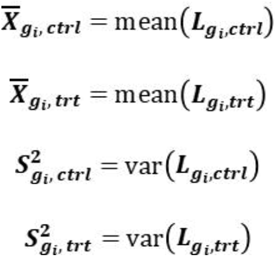

(3) Calculate the input z-score **Z** by the following equation:

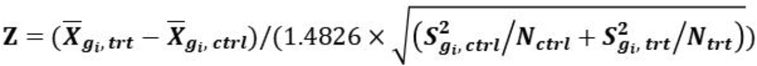

***N*_*ctrl*_, *N*_*trt*_** are the sample sizes of control and compound treated groups respectively. 1.4826 is a scale factor used in LINCS data.

